# Brain-localized CD4 and CD8 T cells perform correlated random walks and not Levy walks

**DOI:** 10.1101/2023.01.09.523257

**Authors:** Dhruv Patel, Raymond Lin, Barun Majumder, Vitaly V. Ganusov

## Abstract

For survival of the organism T cells must efficiently control pathogens invading different peripheral tissues but whether such control (and lack of thereof) is achieved by utilizing different movement strategies remains poorly understood. Liver-localized CD8 T cells perform correlated random walks — a type of a Brownian walk – in liver sinusoids but in some condition these T cells may also perform Levy flights – rapid and large displacements by floating with the blood flow. CD8 T cells in lymph nodes or skin also undergo Brownian walks. A recent study suggested that brain-localized CD8 T cells, specific to *Toxoplasma gondii*, perform generalized Levy walks – a walk type in which T cells alternate pausing and displacing long distances — which may indicate that brain is a unique organ where T cells exhibit movement strategies different from other tissues. We quantified movement patterns of brain-localized *Plasmodium berghei*-specific CD4 and CD8 T cells by using well-established statistical and computational methods. Interestingly, we found that T cells change their movement pattern with time since infection and that CD4 T cells move faster and turn less than CD8 T cells. Importantly, both CD4 and CD8 T cells move in the brain by correlated random walks without long displacements challenging previous observations. We have also re-analyzed the movement data of brain-localized CD8 T cells in *T. gondii* -infected mice and found no evidence of Levy walks. We hypothesize that the previous conclusion of Levy walks of *T. gondii*-specific CD8 T cells in the brain was reached due to missing time-frames in the data that create an impression of large displacements between assumed-to-be-sequential movements. Taken together, our results suggests that movement strategies of CD8 T cells are largely similar between LNs, liver, and the brain and consistent with correlated random walks and not Levy walks.

## Introduction

Different agents starting with large animals such as tigers and wolves and ending with individual cells such as T lymphocytes typically search for resources/targets. Theoretically, different search strategies may have different efficacies depending on the energy used for search, the number and distribution of targets in the environment, the effective dimension of the environment and other details [1]. Finding that a particular type of agent, e.g., T cells, undergo a walk type that is highly efficient in some respects (e.g., in time to find a target) may indicate that such walk types may be evolutionary selected [2]. However, whether observed movement patterns are driven by agent-intrinsic programs or are the consequences of constraints in the environment has rarely been investigated [1, 3–5].

Different search strategies can be roughly subdivided into two large classes: based on Brownian walks and based on Levy walks [6]. The key feature of Brownian walkers is that their displacements are short, consistent with a thin-tailed distribution that has a finite mean & variance [6]. Pure Brownian walkers typically exhibit mean squared displacement (MSD) that changes linearly with time [6–8]. In actual data, agents rarely show MSD to change linearly with time; it is more typical to observe super-diffusion at short time scales due to correlation between sequential speed vectors (so called correlated random walks, CRWs) and sub-diffiusion at longer time scales due to environmental constraints [5, 8–10]. In contrast, Levy walkers typically perform both short and long displacements often consistent with a heavy-tailed distribution composed of a finite mean & infinite variance or both an infinite mean & infinite variance [6]. Levy walkers exhibit super-diffusive behavior that is their MSD increases faster than linear with time [6, 8]. Realistically, however, superdiffusion of Levy walkers may not be always observed over long periods of time due to environmental constraints and other experimental limitations. Both Brownian and Levy walk strategies may have multiple variations; for example, generalized Levy walks are a type of Levy walk in which there are both pauses and runs [2].

Unfortunately, there is no unique, well-established methodology that allows to fully characterize movement strategies of different agents. MSD is often used to determine if agents exhibit normal diffusion (Brownian) or super-diffusion (Levy); however, there are limitations of this method to infer movement types [2, 8–15]. An alternative method is to analyze the distribution of movement/displacement lengths and determine how quickly the tail of the cumulative distribution declines with larger movement lengths [6, 16]. Finally, the distribution of turning angles made by the agents with every displacement can inform about specific movement strategies [7, 17]. Using these methods, many studies including those in immunology have reported movement patterns consistent with Levy walks [2, 18–24]. While it has been common to attribute specific movement patterns to agentintrinsic programs, the relative contribution of internal and external processes in determining why agents move in a specific manner remains typically unknown. For example, we have recently shown that movement patterns of activated CD8 T cells in murine livers such as crawling or floating can be well explained by the physiological aspects of the liver blood vessels (sinusoids) and blood flow [5].

Intravital microscopy is typically used to record movement of various types of immune cells such as T cells or neutrophils [25]. In such imaging experiments, cells to be imaged must express fluorescent markers (e.g., green fluorescent protein, GFP). A small area of the tissue is then scanned by a microscope; a volume of 500 *×* 500 *×* 50 *µ*m is typically scanned every 30 sec. The generated movies are then processed (segmented) by specialized software to determine individual cells and their positions over time. While the movies are typically very impressive at the details of how immune cells behave in tissues in vivo, the resulting coordinate data have many limitations. In particular, 1) cell positions are recorded only at specific time points while cell movements are often continuous; 2) cells may come into or leave the imaging volume over time reducing amount of data available for each cell; 3) segmentation programs may allow cells temporarily to leave imaging volume and introduce missing time frames into the data for individual cell trajectories; 4) while cells are typically moving 3D imaging sometimes only records cell positions in 2D that may bias the interpretation. Exposure tissues to the laser may also impact tissue physiology which in turn may introduce artifacts into the cell position data.

One influential study suggested that brain-localized activated CD8 T cells, specific to *Toxoplasma gondii*, perform generalized Levy walks allowing these T cells to efficiently locate and eliminate *T. gondii*-infected cells [2]. This movement pattern was different from movements T cells exhibit in other tissues such as lymph nodes or the liver suggesting that brain may be a special organ allowing for a unique movement strategy [5, 26]. In this paper we processed recently generated imaging data on brain-localized CD4 and CD8 T cells, specific to *Plasmodium berghei* [27], and characterized movement patterns of these cells using several alternative methodologies. Interestingly, we found that these T cells undergo correlated random walks that are characterized by a relatively short persistence time (*∼* 5 min) and high speeds (7-10 um/min) that are similar to activated T cells localized to the liver. Importantly, brain-localized T cells undergo relatively small displacements that are fully consistent with Brownian walks and are not consistent with generalized Levy walks. We also analyzed the coordinate data of *T. gondii*-specific T cells from Harris *et al*. [2] and found that the data contained missing time-frames for several cells that unless accounted for creates an impression of rare long displacements. After cleaning the data, main characteristics of the movement of *T. gondii*- and *P. berghei*-specific CD8 T cells such as MSD plots and displacement distribution were nearly identical. Our results thus suggest that movement patterns of activated CD8 T cells in the brain are similar to that of the liver and are consistent with correlated (Brownian-like) random walks.

## Materials and methods

### Data

#### Movement data

There are several different datasets used in our analyses. We obtained movies of previously published movements of CD4 and CD8 T cells, specific to *Plasmodium berghei*, in the brains of *Plasmodium*-infected mice [27], which were then segmented using Imaris (Bitplane). We also re-analyzed previously published data [2], provided by Dr. Chris Hunter to VVG. This dataset focuses on movement of activated CD8 T cells, specific to *Toxoplasma gondii*, in the brains of *T. gondii* -infected mice.

#### Imaging Plasmodium-specific CD4 and CD8 T cells in the brain

Imaging was performed in a previous study [27]. Female C57BL/6 (B6) and the transgenic strains PbT-I and PbT-II mice along with the rodent malaria line *Plasmodium berghei ANKA* (PbA) clone 15cy1 were used in this study. The mice were injected i.v. with purified naive PbT-I (CD8) and PbT-II (CD4) T cells (5 × 10^4^) in 0.2 ml PBS. One day later, the mice were infected i.v. with 10^4^ PbA infected red blood cells (iRBCs). For the data used in this experiment, the brains were imaged 6.5 or 7 days post-infection (**Figure 1A**).

**Figure 1:**
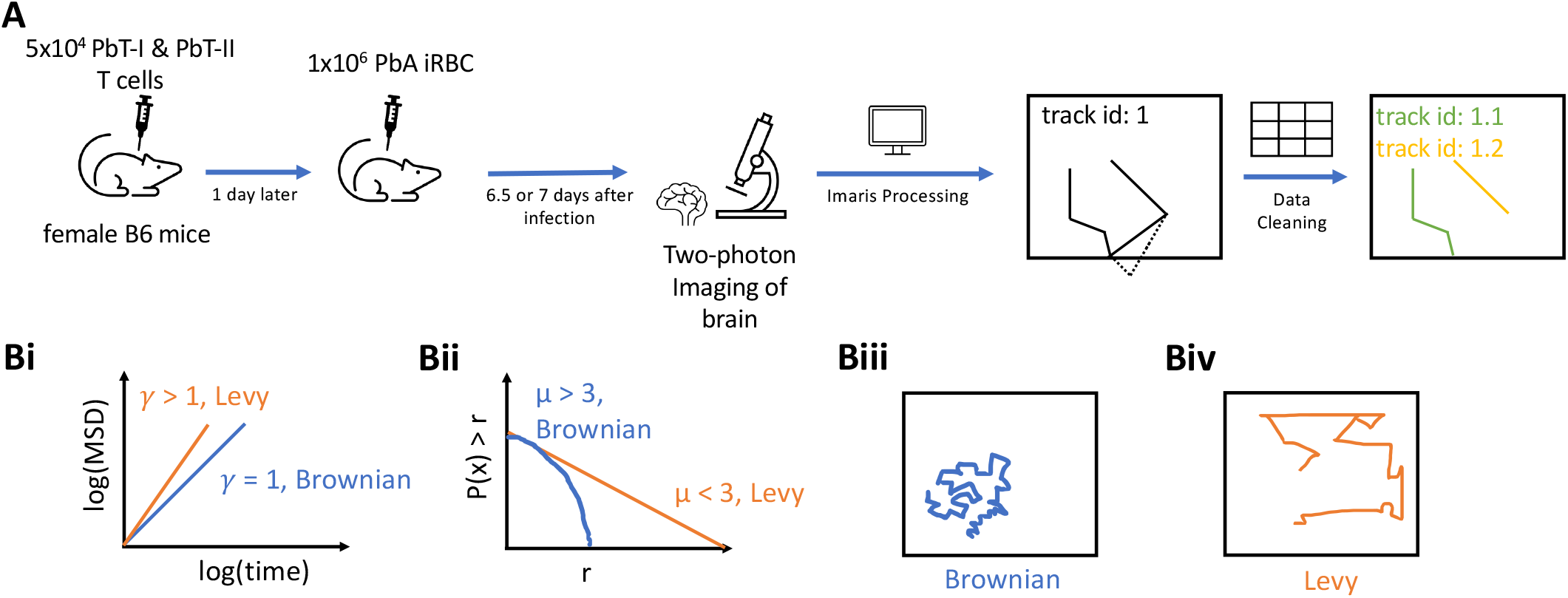
Experimental design and steps in analysis of movement data. **A**: 5 × 10^4^ CD8+ (PbT-I) and CD4+ (PbT-II) T cells were injected i.v. into female B6 mice. Twenty four hours later, the mice were infected i.v. with 10^4^ Plasmodium berghei ANKA (PbA)-infected red blood cells (iRBCs). Six and a half or 7 days post-infection, brains of infected mice were surgically exposed and imaged using two photon microscope at approximately 30 second intervals. Five movies were then processed in Imaris to trace coordinates of CD4 and CD8 T cells. Any tracks that contained gaps in time were split. **B**: To characterize cell movement we calculated the mean square displacement (MSD, Bi) and the movement length distribution (Bii); estimated parameters *γ* and *µ* allowed to determine if the cells exhibit Brownian (Biii) or Levy (Biv) walks.

#### Processing movies from PbA-infected mice with Imaris

We analyzed a total of five movies from Ghazanfari *et al*. [27] experiments (**Figure 1A**). Within each movie there were multiple channels; however, only the channels containing signal of the CD8 T cells (PbT-I) and CD4 T cells (PbT-II) were relevant to our analysis, and these were the channels we used for the analysis. We used function “Spots” in Imaris (https://imaris.oxinst.com/) to identify and track individual T cells. Depending on the cell parameters chosen in Imaris, it is typical for Imaris’ basic algorithm to identify many more objects that there may be in reality. Every movie was manually inspected and objects that were not likely correspond to T cells were removed. We also removed objects that may result in biased interpretation including i) T cells being close to the imaging border, ii) cells were sliced in half, or iii) immobile cells. By visually inspecting each movie and each track, we manually joined tracks of cells that were incorrectly split into independent tracks by Imaris. We also manually traced T cells that were clearly visible but missed by the standard algorithm of Imaris. Additional details of data segmentation are shown in Supplemental Information.

#### Cleaning cell position data

When tracking cells, default settings of Imaris allow several positions of a cell to be missing and still be considered as a single track. Unless correct, such segmentation results in missing time-frames in the cell coordinate data that may result in misspecification of cell movement types. All of our datasets contained cell trajectories with missing time frames. Following our previously outlined methodology, we “cleaned” the cell position data as follows (**Figure 1A** and [5]):

1. Checked for duplicates in the data and assign unique track IDs to each of them. We used trailing letters in this case.
2. For any time gap between tracks of the same track ID that is greater than the imaging frequency plus a certain offset (1 second in our case), the track for that track ID should be split into two tracks with unique track IDs at that point. This is iterated until there are no time gaps larger than the imaging frequency plus the offset between tracks of the same track ID.
3. All tracks are then shifted to begin at t = 0 sec.

We initially had the data for 264 PbA-specific CD4 T cells; we generated 149 new (split) trajectories resulting in 413 T cells. After removing trajectories with only a single position we were left with data for 355 tracks with 5247 positions. Similarly, for the total of 589 PbA-specific CD8 T cells we generated 458 new trajectories resulting in 1047 T cells. After removing trajectories with only a single position we were left with data for 873 tracks with total 10563 positions.

#### Time standardization

Intravital imaging movies were generated with slightly different imaging frequencies: 30 sec, 30.68 sec, & 32.76 sec. To combine all track data into one dataset we assumed that all movements were done in 30 sec time intervals.

#### Published dataset on movement of *Toxoplasma gondii* -specific CD8 T cells in the brain

We obtained the data from Harris *et al*. [2] on movement of brain-localized CD8 T cells, specific to *T. gondii*. The data consisted of two separate datasets with different imaging frequencies (19.25 sec & 22 sec). To increase the power of analysis, we merged the datasets into one with the assumed imaging frequency of 20 sec. These data also had many cell trajectories that had missing time-frames that were split to ensure consistent time intervals between sequential cell movements. We initially had the data for 657 CD8 T cells; we generated 217 new cells resulting in a total of 874 T cell trajectories. After removing tracks with only a single position we were left with data for 812 T cells with 18848 positions.

#### Statistical analyses

To characterize movement type of T cells, we followed a methodology outlined in our previous pub-lication [5].

#### Arrest coefficient and meandering index

We calculated the arrest coefficient which is the percent of the time each T cell was arrested (instantaneous speed *<* 2 *µ*m/min) [28]. We also computed the meandering index which is the straightness of each T cell trajectory defined as the ratio between the overall displacement and the total distance traveled [29].

#### Mean square displacement

We also calculated the mean square displacement (MSD). While MSD can be calculated for individual cells or for the whole population (e.g., [30]), here we used all tracks in a given dataset where imaging frequency was either the same or standardized to calculate the MSD using formula

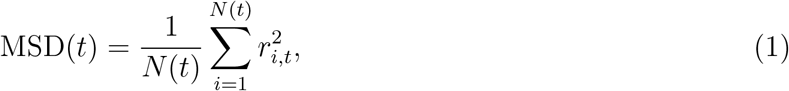

where *r*_*i,t*_ is the displacement of an *i*^*th*^ cell from its initial position to the position at time *t* and *N* (*t*) is the number of cells for which coordinate data was available at time *t*. To characterize the MSD, we used the relationship MSD(*t*) = *ct*^*γ*^ where *t* is time delay and *γ* was estimated by linear regression via log-log transforming MSD and *t* for a subset of data that lies on the regression line. The estimate of *γ* can be used to characterize cell movement type (**Figure 1Bi**): *γ* < 1 suggests subdiffusion, indicating that cells are constrained spatially in movement, *γ* = 1 suggests that cells demonstrate Brownian movement, and *γ* > 1 suggests superdiffusion of cells, indicating that cells demonstrate directed movement or transportation [8]. It should be noted, however, that some types of Brownian walks, for example, correlated random walks may display transient superdiffusion [11].

#### Basic distributions to characterize cell movements

To characterize the type of walks performed by agents, a number of different distribution has been used. Our recent work suggested that the generalized Pareto distribution (GP) fits to movement length distribution of liver-localized CD8 T cells with the best quality [5]. An interesting property of the GP distribution is that it can describe both Brownian and Levy walks and can be fitted to the whole dataset on movement length distribution [5]. The GP distribution is defined as

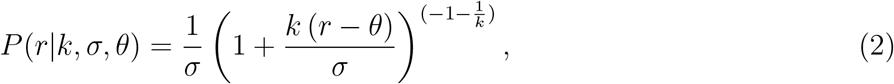

where *k* is the shape parameter, *σ* is the scale parameter, and *θ* is the location parameter. The mean of this distribution is 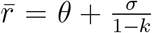 and variance is 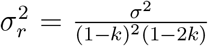. When *k >* 0 and *θ* = *σ/k*, GP simplifies to the Pareto (or powerlaw) distribution

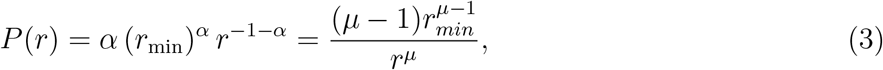

where *r* ≥ *r*_min_ is the displacement, 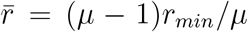, and *r*_min_ = *σ/k* is the scale parameter; *α* = 1/*k* and *µ* = *α* + 1 are the shape parameter. The Pareto distribution typically cannot fully describe the distribution of movement lengths of T cells, so it is fitted to the tail of the data that includes a subset of longest displacements [16]. Such “tail analysis” (see below) allows to estimate the shape parameter *µ* that is used to characterize the walk type as Brownian (*µ* > 3) or Levy (*µ* < 3, [16]). Cases with *µ* < 2 corresponds to a subset of Levy walks called bullet motion [6].

#### Fitting GP distribution to data

We fit the models (eqn. (2)) to distribution of movement lengths *r* of cells calculated as the distance traveled by each cell between two sequential time points. To make sure that these displacements are calculated for the same time intervals, data were cleaned by splitting the trajectories (see above). The likelihood of the model parameters given the data is given as

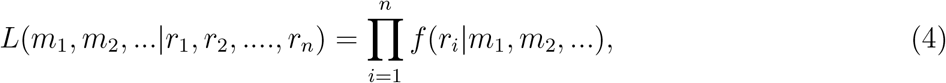

where *r*_*i*_ are the cell movement length data consisting of *n* data points. The probability density function *f* (*r*_*i*_|*m*_1_, *m*_2_, …) is given in eqn. (2) (or can be any other distribution, e.g., eqn. (3)). Parameters *m*_1_, *m*_2_, … of the models were estimated by minimizing the log of negative likelihood *ℒ* = *−* ln *L*. Fitting the model to data was done in python using function genpareto.fit in package SciPy.

#### Tail analysis of movement length distribution

In addition to fitting the GP distribution to the whole dataset of cell movement lengths, we used a previously suggested tail analysis [16, **Figure 1Bii**]. To fit the Pareto distribution to the subset of the data it is critical to define which movement lengths will remain the data and which will be ignored, i.e., to define *r*_min_ in the Pareto distribution (eqn. (3)). The methodology of the rigorous approach to define *r*_min_ has been outlined previously [16] and implemented in the python package *powerlaw* [31] that we used. The package takes an input of movement lengths and outputs the estimate of *r*_min_ and shape parameters *µ*. In some cases, parameter *r*_min_ could be defined by the user and not estimated from the data. This package uses the likelihood approach (eqn. (4)) to fit the Pareto distribution to the movement lengths data for different values of *r*_min_ and compares the quality of the model fit to data using a statistical test [16].

#### Comparing the data

Unless otherwise noted we used Mann-Whitney tests to compare average/median values for two samples, e.g., difference between average meandering index for CD4 vs. CD8 T cells.

### Simulations

To investigate whether the previously proposed methodology of determining the type of cell walk by fitting GP distribution to all movement length data or by fitting the Pareto distribution to the subset (tail) of the data are appropriate we simulated cell movements as Brownian or Levy walkers where the parameters of cell movement are known [5, 16, 31]. To simulate movement of Brownian or Levy walkers we generated random numbers of cell movement lengths from a Pareto distribution (eqn. (3)) with parameter *µ* for every time step. In the case of Brownian walkers we chose *µ* > 3 and in the case of Levy walkers *µ* < 3; specific value of the shape parameter *µ* was varied in different simulations, and we used a default value for *r*_min_ = 1. We simulated movements in 3D by using von Mises-Fischer (vMF) distribution to generate random 3D vectors with the concentration parameter *κ* = 0.01 ensuring no preferred direction for cell movement [17, 32]. To simulate T cell movement as generalized Levy walkers we used the methodology outlined previously [2]. Specifically, we created two Pareto distributions (eqn. (3)) with two shape parameters: *µ*_*run*_ *<* 3 and *µ*_*pause*_ *<* 3. Since generalized Levy walkers can pause after each run, we incorporated a pause after each displacement. A pause time would be selected from the *µ*_*pause*_ of the Pareto distribution and pause time was a discrete number of time steps used in simulations. After the pause time elapses, the cell will perform a movement. If the cell was to move rather than pause, a displacement length would be selected from the *µ*_*run*_ Pareto distribution, and the cell would then proceed to pause again. The direction of displacement is selected from a vMF distribution with *κ* = 0.01. All simulations were done in python. Random numbers of the Pareto distributions were generated using the python package *SciPy*.

## Results

### Experimental design and modeling analyses to rigorously characterize movement pattern of brain-localized T cells

T cells are typically activated in secondary lymphoid organs, and following activation and differentiation migrate to peripheral tissues to control infections. Whether T cells utilize specific, evolutionary-selected strategies, to locate and eliminate infections in peripheral tissues remains debated. Brain is an immunopriveledged site, and movement strategies of brain-localized T cells may be unique [2]. Ghazanfari *et al*. [27] developed novel TCR transgenic mice with T cells recognizing antigens of Plasmodium berghei ANKA (PbA) on MHC-I and MHC-II molecules, allowing to track PbA-specific CD8 and CD4 T cell response with intravital imaging (**Figure 1**). In these experiments, mice received fluorescently labeled naive PbA-specific T cells, PbA-infected RBCs, and the movement of the labeled T cells in murine brains was followed with intravital microscopy. We then processed the imaging data with Imaris and cleaned the resulting cell trajectory data to avoid instances where cell positions were not clearly defined (**Figure 1A** and see Materials and methods for more detail). In total, data from 5 movies were analyzed: 3 movies for 6.5 days post infection and 2 movies for 7 days post infection. We then applied a number of statistical methods to rigorously characterize movement pattern of T cells and whether these movements are consistent with Brownian or Levy walks (**Figure 1B**).

From our imaging analyses we found that in the imaging area of 512 × 512 × 44 *µ*m there were more PbA-specific CD8 T cells than CD4 T cells, and while the number of CD4 T cells slightly declined with time (84 vs. 81 cells/movie for 6.5 vs. 7 days post-infection), the number of CD8 T cells increased with time since infection (145 vs. 305 cells/movie, **Figure 2A** and **Supplemental Movie S1**); both changes, however, were not statistically significant due to a small number of movies analyzed. The constancy/decrease in CD4 T cell numbers is not fully consistent with flow cytometry data on kinetics of brain-localized endogenous T cell response to PbA [27]. Given the number of cells observed in these imaging experiments, we next calculated the potential number of brain-localized T cells after PbA infection. The total volume imaged in these experiments was 512 × 512 × 44 *µ*m^3^ = 0.0115 mm^3^, and given the volume of the brain of B6 mice of is 509 mm^3^ [33], the scaling factor to calculate the total number of cells per brain given the number of cells in the imaging area is 509*/*0.0115 = 4.42 × 10^4^. This translates to 3.71 × 10^6^ and 3.55 × 10^6^ PbA-specific CD4 T cells at 6.5 and 7 days post-infection, and 6.38 × 10^6^ and 1.35 × 10^7^ PbA-specific CD8 T cells at 6.5 and 7 days post-infection. Interestingly, 100-fold lower numbers were calculated for brain-localized T cells obtained using flow cytometry [27].

**Figure 2:**
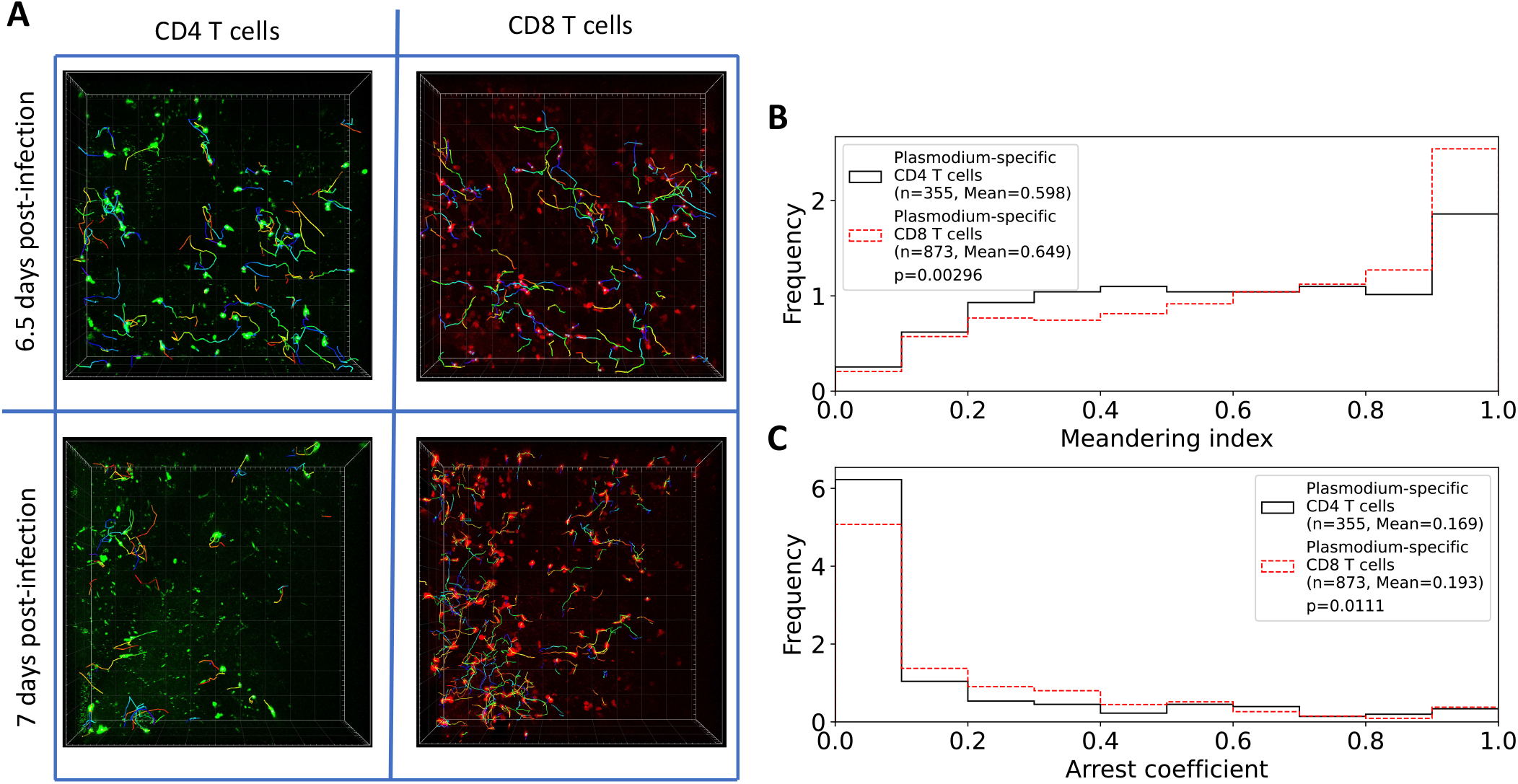
Brain-localized *Plasmodium* -specific CD4 and CD8 T cells differ in kinetics of accumulation in the tissue and movement. **A**: Following experimental design (**Figure 1**) we segmented 3 movies for 6.5 days and 2 movies for 7 days since PbA infection and calculated positions of brain-localized CD4 and CD8 T cells **(A)**. Tracking of T cells was done using Imaris (see Materials and Methods for more detail). In total we analyzed tracks for *n* = 252 CD4 and *n* = 434 CD8 T cells at 6.5 days post infection, and for *n* = 161 CD4 and and *n* = 613 CD8 T cells at 7 days post infection. **B**: meandering index for all CD4 and CD8 T cells. **C**: arrest coefficient for all CD4 and CD8 T cells. Examples of the movies are shown in **Supplemental Movie S1**. Statistical comparison in panels B-C was done using Mann-Whitney test with *p*-values from the test indicated on individual panels.

In addition to the difference in the number between CD4 and CD8 T cells we found that their overall movement patterns were also slightly different: CD4 T cells have lower meandering index than CD8 T cells indicating less straight paths between sequential time frames (0.598 vs. 0.649, *p* = 0.003); CD4 T cells also spend less time pausing (speeds < 2 *µ*m/min) with a slightly lower arrest coefficient (0.169 vs. 0.193, *p* = 0.01, **Figure 2B-C**).

### CD4 and CD8 T cells in the brain undergo correlated random walks

A previous pioneering study proposed that brain-localized CD8 T cells perform generalized Levy walks characterized by rare but long displacements [2, **Figure 1Biv**]. Visual inspection of the five movies did not reveal any long displacements by T cells; however, there were some visible cell-like objects moving rapidly with the blood flow similar to what we have observed for liver-localized CD8 T cells [5]. We therefore performed rigorous analyses of the cell trajectory data by pooling data from different experiments together. An important step in the analysis was our recent recognition of missing time-frames in cell position data generated by Imaris and providing methodology to “clean” such data (see Materials and methods for more detail) [5].

We first calculated the mean squared displacement (MSD) for brain-localized *Plasmoidum*-specific CD4 and CD8 T cell trajectories (eqn. (1)). Both cell populations displayed super-diffusive MSD that increased faster than linearly with time (slopes *γ* = 1.59 and *γ* = 1.39, for CD4 and CD8 T cells, respectively, **Figure 3A**). The superdiffusion was transient for about 5-7 min as MSD saturated at longer times as expected. Interestingly, fitting Fürth equation to the MSD plots suggested much shorter persistence time of T cells of 1 min [34, results not shown]. While superdiffusion is the main feature of Levy walkers [8], some types of Brownian walks such correlated random walks can also display superdiffusion [11].

**Figure 3:**
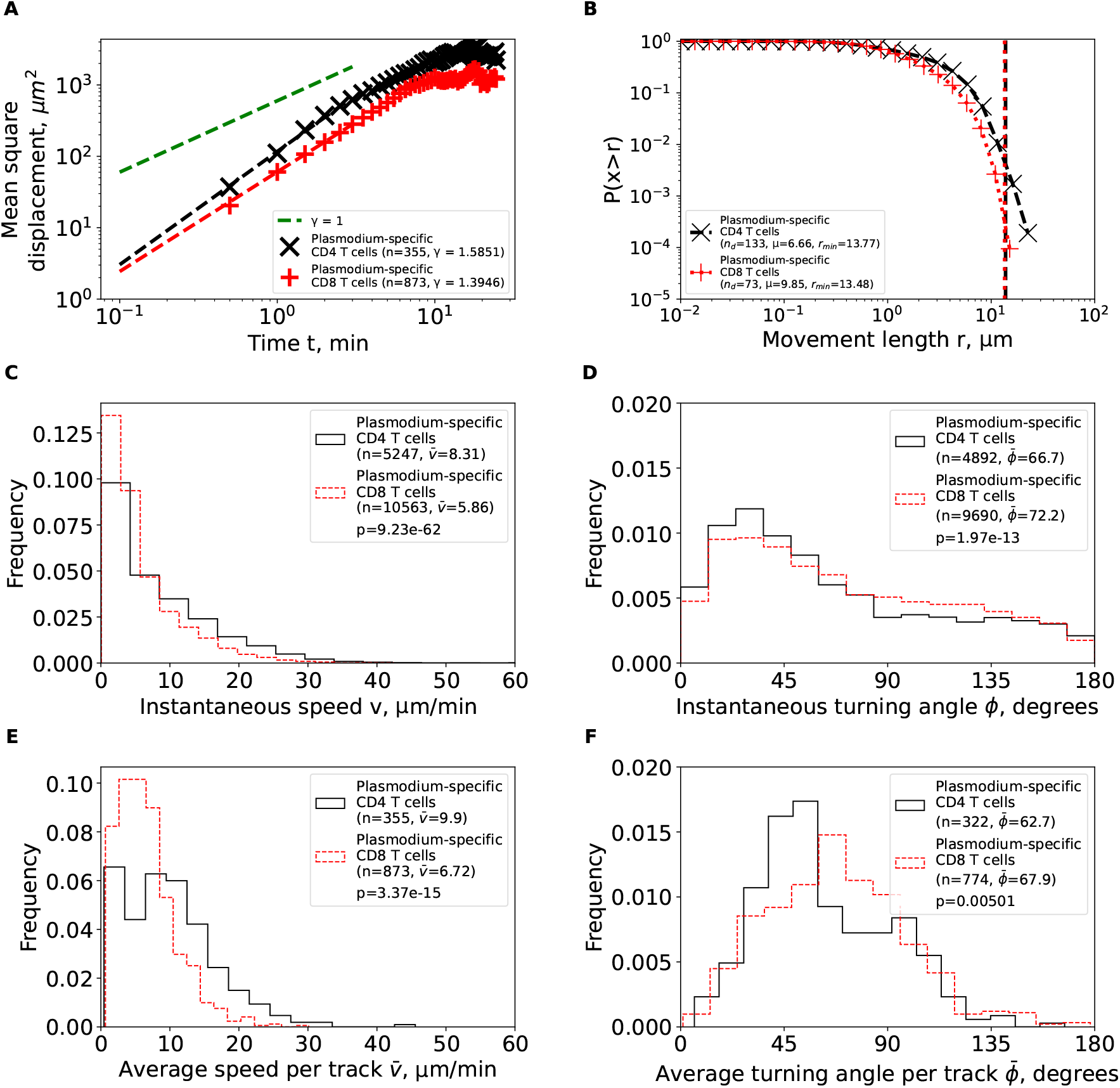
Brain-localized *Plasmodium* -specific CD4 and CD8 T cells undergo correlated random walks with transient super-diffusive displacement. We calculated basic movement characteristics for CD4 and CD8 T cells in brains of PbA-infected cells (**Figure 1**) such as mean squared displacement (MSD, **A**), movement length distribution (**B**), instantaneous speeds (**C**) and turning angles (**D**), and average speed (**E**) and turning angle (**F**) for each track. The slope of the linear regression for the log(MSD) with respect to log(*t*) for first several minutes is denoted by *γ*. Estimated parameters of the Pareto distribution (eqn. (3)) fitted to the tails of the step length distribution are denoted as *µ* and *r*_min_ (see Materials and Methods for details). Difference in averages/medians was calculated using Mann-Whitney tests with associated *p*-values from the test indicated on individual panels (C-F). All panels indicate the number of measurements/data points *n* used in analysis and the average values for the parameters. Note that all calculations were performed on cleaned T cell tracks in which tracks with missing time frames were split into separate tracks (**Figure 1A** and see Materials and methods for more detail).

We then analyzed the cumulative distribution of movement lengths which was similar for CD4 and CD8 T cells; interestingly, there were no very long displacements in these data (**Figure 3B**). By using an established “tail” analysis that fits the Pareto distribution to the tail of the cumulative movement length distribution (i.e., to a subset of data with longest movements) we found shape parameters for CD4 T cells (*µ* = 6.66) and CD8 T cells (*µ* = 9.85) to be consistent with Brownian (*µ* > 3) and not Levy (*µ* < 3) walks (**Figure 3B**). To ensure that data subselection in the tail analysis did not bias the results we fitted the generalized Pareto (GP) distribution (eqn. (2)) to all data using a likelihood approach (eqn. (4)); we have previously found that GP distribution provides the best fit of the movement lengths data for liver-localized CD8 T cells [5]. Importantly, the GP distribution fitted the data well and predicted a finite mean and variance for data for both T cell populations (**Supplemental Table S1**). A finite mean and variance are features of Brownian movement and not of Levy walkers [6].

We finally investigated if other cell movement characteristics may be inconsistent with correlated random walks. Distribution of instantaneous speeds suggested that no cell had extraordinary speeds with maximum not exceeding 40 *µ*m/min (**Figure 3C**). Turning angle distribution for both cell types was biased towards acute angles which is a key feature of correlated random walks (**Figure 3D**). On average, CD4 T cells had higher instantaneous speeds than CD8 T cells 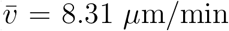. 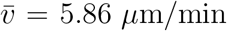, **Figure 3C**) and smaller turning angles 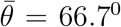. 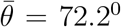, **Figure 3D**). There were more CD4 T cells displaying high speeds (> 20 *µ*m/min) than CD8 T cells (**Figure 3E**), and CD4 T cells exhibited lower average turning angles (and thus higher persistence in movement in one direction) as compared to CD8 T cells 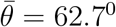 vs. 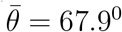. Taken together, our analyses suggest that *Plasmodium*-specific, brain-localized CD4 and CD8 T cells performed correlated random walks with relatively high speeds and low turning angles and displayed transient super-diffusive displacement.

### T cells show increased speeds and turning as time after infection increases

For our first set of analyses we pooled the trajectory data from imaging performed at 6.5 or 7 days after infection. We next sought to determine if movement patterns were different between the two times since infection. Importantly, regardless of time after infection, all cell populations exhibited features of correlated random walks with transient superdiffusion (*γ* > 1 in **Supplemental Figure S1A** and **Supplemental Figure S2A**) and relatively short movement lengths (*µ* > 3 in **Supplemental Figure S1B** and **Supplemental Figure S2B**). Additionally, best fits of the GP distribution predicted a finite mean and variance further supporting that these cells exhibited Brownian-like walks [6].

Time since infection had a different impact on other movement characteristics of T cells. CD4 T cells increased their instantaneous speeds and turning angles between 6.5 and 7 days post infection (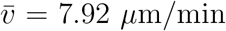 vs. 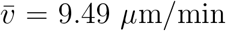 and 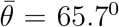 vs. 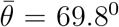, respectively, **Supplemental Figure S1C-D**). Interestingly, however, the average speed or average turning angle per cell did not significantly change between 6.5 to 7 days post-infection (**Supplemental Figure S1E-F**). In contrast, CD8 T cells increased their instantaneous and average per cell speeds between 6.5 and 7 days post-infection 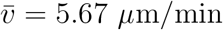 vs. 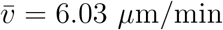 and 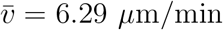 vs. 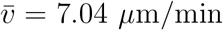, **Supplemental Figure S2C**&E) and the instantaneous and average (per cell) turning angles also increased with time since infection 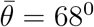 vs. 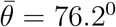 and 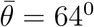 vs. 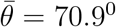, **Supplemental Figure S2D**&F). The overall (although inconsistent) increase in speed and turning angle of both CD4 and CD8 T cells with time since infection may be related to the need for T cells increase efficiency of search for pathogen at later times after infection [35–37].

### Uncleaned trajectory data may result in misspecification of the walk type

In our analyses we found that brain-localized CD4 and CD8 T cells perform Brownian-type (correlated random) walks which contradicts a previous result suggesting that movement patterns of CD8 T cells in the brain are consistent with generalized Levy walks [2]. We obtained the coordinate data of brain-localized CD8 T cells from Harris *et al*. [2] and carefully inspected these data. We found that there were several technical issues with the data. First, the data had many missing time-frames (*n*_*gaps*_ = 217, **Supplemental Figure S3A**) which, if not cleaned, can produce long displacements between two-assumed-to-be-sequential time-frames. Several of these gaps are substantial (6 and 16 time-steps, **Supplemental Figure S3A**). Second, the data contained a duplicate name tag for a track that eventually diverged. This resulted in an extremely long displacement (80.4 *µ*m, **Supplemental Figure S3B**) in the span of one time-frame (22 seconds). We performed tail analysis on the movement length distribution data and found that the best fit is found with the shape parameter *µ* = 3.17 which is consistent with Brownian and not Levy walks (**Supplemental Figure S3C**). However, by varying the scale parameter *r*_min_ we did find instances when shape parameter *µ* < 3 suggesting that these data may be consistent with Levy walks (**Supplemental Figure S3C**). However, after cleaning the trajectory data (by splitting trajectories that have missing time frames), tail analysis resulted in highly consistent estimates of the shape parameter *µ* > 3 for all reasonable values of *r*_min_ > 5 *µ*m consistent with Brownian walks (**Supplemental Figure S3D**).

### *Plasmodium* - and *Toxoplasma* -specific, brain-localized CD8 T cells behave similarly

To more rigorously characterize movement pattern of *Toxoplasma*-specific, brain-localized CD8 T cells from Harris *et al*. [2] study we performed the same analyses as we did for *Plasmodium*-specific T cells. Similar to *Plasmodium*-specific CD8 T cells, the *Toxoplasma*-specific CD8 T cells also displayed super-diffusive displacement (*γ* = 1.54, **Figure 4A**), with rapidly declining distribution of movement lengths (*µ* = 5.34, **Figure 4B**). Fits of the GP distribution to the movement length distribution data also predicted finite mean and variance (**Supplemental Table S1**). Despite targeting different parasites, other movement parameters were quite similar between *Toxoplasma*- and *Plasmodium*-specific CD8 T cells including similar instantaneous and average (per cell) speeds and turning angles (**Figure 4C-F**). Taken together, our analysis suggests that movement patterns of brain-localized CD8 T cells are similar between two infections and two independent studies, and are consistent with Brownian rather than Levy walks.

**Figure 4:**
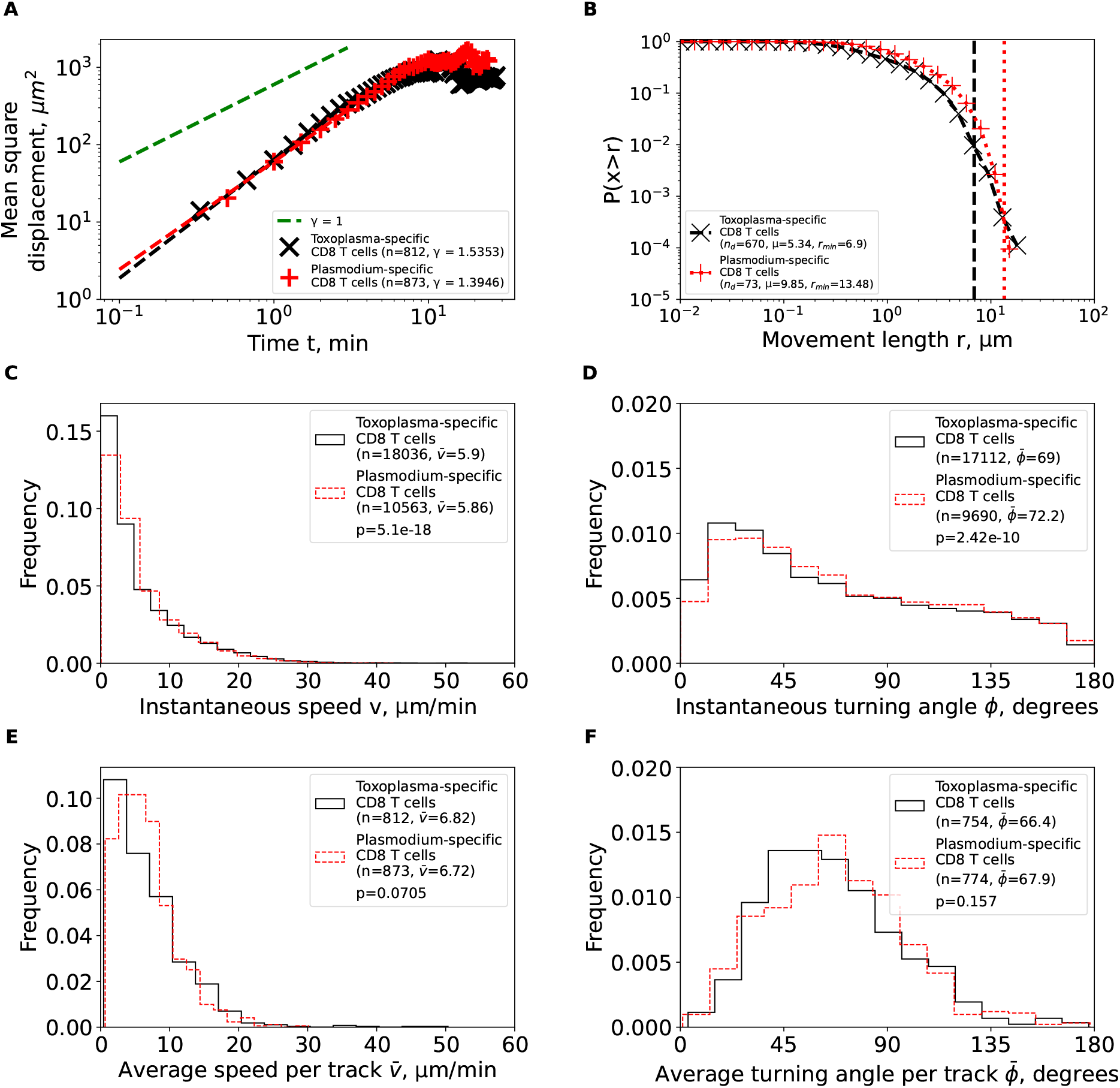
Brain-localized CD8 T cells specific to *Plasmodium berghei* or *Toxoplasma gondii* behave similarly as Brownian and not Levy walkers. We calculated the same characteristics as in **Figure 3** for *Toxoplasma gondii*-specific CD8 T cells from the previous publication [2] after cleaning and renaming the cell tracks (see Materials and methods for more detail). For comparison, the estimated characteristics for PbA-specific CD8 T cells from **Figure 3** are plotted on the same graph to illustrate similarities in movement characteristics.

### Our methods of determining walk type are robust

While we have used previously proposed methods to determine the type of cell walk [16, 38], there may be a possibility that these methods may fail for our trajectory data. Therefore, to check validity of these methods we simulated cell movements as Levy walkers, generalized Levy walkers, and bullet motion walkers and fitted the GP distribution to these data (see Materials and Methods for details on simulations). In cases of Levy walkers we found that variance of the best fit GP distribution is infinite which is consistent with Levy walks [6]. However, in all cases when simulated movements were non-Brownian we found that GP fits indicated infinite variance (and sometimes mean) suggesting that this approach allow to detect deviations from Brownian walks. Additionally, we also determined the walk type by fitting the Pareto distribution to the tail of the movement length data for simulated generalized Levy walkers when using a large range of possible parameters. We found that the tail analysis accurately determined the shape parameter *µ* (**Supplemental Figure S4**). We only found few exceptions when we used *µ* ≈ 3 but found *µ* > 3 in the fits which we believe is likely due to a combination of chance and package sensitivity.

## Discussion

One could expect that T cells should utilize different movement strategies when responding to diverse pathogens and migrating to different lymphoid or non-lymphoid (peripheral) tissues. Brain is an important immunopriveledged site, and one recent study suggested that brain-localized CD8 T cells search for infection using generalized Levy walks that may increase the search efficiency, thus, may potentially minimize bystander damage to the delicate brain tissue [2]. Here we rigorously analyzed data from novel experiments imaging movement of *Plasmodium*-specific CD4 and CD8 T cells in brains of PbA-infected mice (**Figure 1**). By using alternative complementary tools we found that brain-localized T cells display transient superdiffusive displacement which is achieved by making small movements in the same direction (i.e., small turning angles) — both are cardinal features of Brownian-like (correlated random) walks (**Figure 2**).

While the overall pattern of T cell movements were consistent with correlated random walks, there were differences between CD4 and CD8 T cells in details of the movement and how the movement was impacted by the time since infection. In particular, CD4 T cells moved faster and turned less than CD8 T cells (**Figure 3**). T cells also had higher speeds and higher turning angles between day 6.5 and 7 days post infection even though specific details slightly varied whether we compared instantaneous or per cell average speeds and turning angles (**Supplemental Figures S1 and S2**). Increasing speeds and turning may allow for a more thorough search of the infection but also could be a stress response. Imaging did allow to detect some parasites in brains of PbA-infected mice [27] suggesting that the parasites are likely to be present in the brain blood vessels and thus may results in mechanical obstruction of blood vessels as observed in human cerebral malaria [35–37]. Obstruction of brain blood vessels can lead to hypoxia [35], which may have caused the T cells to enter a “panic mode” later in infection. This would explain the increase in speed and turning angle with respect to time after infection in same cell types.

By using these imaging data we predicted that brains of PbA-infected mice should contain 3*−* 10 × 10^6^ CD4 or CD8 T cells. However, flow cytometry-based measurements suggest only presence of 0.1 *−* 5 × 10^4^ T cells at these early (6.5 days) times since infection [27]. Calculating the number of T cells in the whole brain from imaging only 0.002% of its total volume may be problematic as T cells tend to leave and enter the imaging area. Using the number of T cells identified per only one time frame did not dramatically change the estimate of the total number of cells in the brain (results not shown). These results are consistent with a previous observation of gross under-estimation of the number of T cells isolated from many nonlymphoid tissues for flow cytometry-based analyses [39]. Because extraction of lymphocytes from nonlymphoid tissues may generate biases (e.g., non-specific enrichment for some cell subtypes, [39]) caution needs to be taken when interpreting differences in numbers and phenotypes of brain-localized T cells from flow cytometry analyses.

Our results that brain-localized, PbA-specific T cells perform correlated random walks are in contrast with results of another study suggesting that brain localized, *Toxoplasma*-specific CD8 T cells perform generalized Levy walks [2]. When we cleaned and reanalyzed Harris *et al*. [2] data we found that movement patterns of PbA- and *Toxoplasma*-specific CD8 T cells are very similar, both displaying transient superdiffusion and short movement lengths (**Figure 4**). We believe that the difference arose in part because of how the original position data were analyzed – we found that missing time-frames for some cell trajectories created unreasonably long movements that provided some support for Levy walks (**Supplemental Figure S3**); when we splitted into these movements into new tracks we found no long movements, and this ultimately allowed us to classify movement of *Toxoplasma*-specific CD8 T cells as Brownian-like walkers. Another limitation of the Harris *et al*. [2] study was relatively poor description of how the models were fitted to movement length data and how the quality of the model fits was evaluated.

Our finding that brain-localized T cells undergo correlated random walks is consistent with finding of a similar movement program of activated CD8 T cells in the liver or skin and naive/resting CD8 T cells in lymph nodes [5, 26, 40]. CD8 T cell movements in other non-lymphoid tissues such as the lung have not been tested if they are consistent with Brownian or Levy walks [41]. Given the ability of T cells in tissues to do a persistent walk in a given direction (cell “inertia” [17]) and inability of T cells to make very long displacements in constrained tissue environments, it is likely that Brownian-like, correlated random walks are the rule for T cell movements in tissues rather than exception. Yet, specific details of these Brownian-like, persistent movements may vary significantly between different tissues including differences in instantaneous and average (per cell) speeds and turning angles. These details may dramatically impact the overall efficiency at which T cells survey the organ, and could be a result of specific sub-populations of T cells entering different tissues or could simply be due to different physiological constraints imposed by the environment. For example, CD8 T cells in the skin have much lower average speeds than CD8 T cells in the liver (3 *−* 4 *µ*m/min vs. 5 *−* 15 *µ*m/min, [5, 40, 42]) although as far as we know none of previous studies have imaged T cells in multiple tissues in the same animal. Other factors such as the degree of inflammation in the tissue and imaging frequency may also influence inferred parameters such as speeds and turning angles [17, 32].

Several recent studies have suggested there may be a cell-intrinsic program that links ability of cells to move and to turn so that cells with higher speeds turn less [14, 43]. We found that indeed, T cells that have higher speeds tend to have lower turning angles (so they turn less); for example, PbA-specific CD4 T cells have higher average speeds and lower average turning angles than PbA-specific CD8 T cells (**Figure 2**). Furthermore, we found a strong negative correlation between average speed and average turning angle per cell for the whole population of CD4 or CD8 T cells (results not shown). We have recently shown that a negative correlation between average speed and average turning angle may simply arise as a sampling artifact [17]. Indeed, we found that between 6.5 and 7 days post PbA infection both speeds and turning angles of T cells increase challenging the idea that the correlation between speed and turning is cell-intrinsic (**Supplemental Figure S2**).

Our study has several limitations. Intravital imaging involved only a small cross-section of the brain which is limited the sample size and creates a bias of generating longer trajectories for slower and more centrally located cells. Our analysis also assumes homogeneity among cells in a population as for most of our analyses we pooled the data from individual movies to increase the statistical power. It may be possible that there are subpopulations of cells that exhibit different movement patterns. Given that in intravital imaging experiments it is typically hard to follow individual T cells for longer than 15-20 time frames, characterizing heterogeneity of movement of individual cells in vivo remains difficult (but see [14]). Missing time frames for some cell trajectories necessitate to “clean” such data (split the tracks) further reducing amount of data available to rigorously quantify heterogeneity in T cell movement behavior in vivo. We found that different metrics used to evaluate how persistent T cells are in their movement may give different answers. For example, CD4 T cells had lower turning angles than CD8 T cells and yet CD8 T cells had larger meandering index – lower turning angles and larger meandering index suggest straighter trajectories (**Figure 2B** and **Figure 3F**). There is a need for rigorous studies that compare these and other metrics such as persistence time to better understand conditions under which these metrics truly characterize ability of T cells to exhibit persistent movement. We only had access to five movies to analyze, and it is possible that collecting more data may reveal deviations of T cell movement patterns from Brownian-like walks. However, it remains hard to justify long displacements of T cells found in some studies (e.g., [2]) given physiological constraints imposed by tissues on moving T cells. One exception to this rule are cells located in vasculature that allows such cells to float with the blood flow exhibiting Levy flights [5]. Physiological relevance of these floating events to the search efficiency remains to be established, however. While imaging frequency used in Ghazanfari *et al*. [27] (30 sec) was sufficient to accurately detect movement of brain-localized T cells, it was clearly insufficient to better understand movement patterns of T cells in the vasculature, and as these cells move from the blood vessels into the parenchymal tissues. Using a higher frequency imaging may be needed to quantify movement patterns of T cells, located in the blood vessels of the brain [42]. Finally, we did not have access to the original imaging data from Harris *et al*. [2] study, and therefore, in our analysis we only relied on the data provided by Harris *et al*. [2] and cannot comment on validity of the data except some technical issues such as missing time-frames for some cell trajectories.

Our results raise questions for future research. There is a need for sanity checks of the coordinate data generated from intravital imaging experiments; in particular, whether such data have missing time frames and thus may require cleaning. We have provided a python-based script for cleaning trajectory data. It may be interesting to extend our calculations of the total number of T cells found from intravital imaging experiments and using more traditional approaches, e.g., flow cytometry, to other tissues and infections. It is possible that many more T cells localize the to peripheral sites that sometimes is assumed and this is likely to provide better estimate of the overall magnitude and kinetics of T cell responses to pathogens [39]. Whether there are true differences in the movement patterns (e.g., speeds, turning angles) between CD4 and CD8 T cells in other infections and tissues may need to be explored. While we know that T cell populations are heterogeneous, we need better tools to track individual T cells from longer periods of time so we can rigorously evaluate the degree of kinetic heterogeneity of these cells in different settings. While we have shown that GP distribution can well describe movement length data of T cells in the liver and the brain, we need to test the same model fits the data best in other cases. Furthermore, we need additional parametric distributions to describe other features of T cell movements, e.g., turning angles. Using vMF distribution may be particularly useful for that purpose [17, 32]. We expect that environmental constraints should influence movement patterns of T cells [5]; thus, there is a need to visualize and quantify these constraints. Finally, basic motility parameters for T cells in various tissues should be used in mathematical models to predict the optimal numbers of T cells needed to prevent or control infections. This would require a close collaboration between experimentalists and modellers and such collaborations should be explicitly supported by funding agencies.

## Abbreviations

GP: generalized Pareto,
PbA: Plasmodium berghei ANKA,
iRBCs: infected red blood cells,
MSD: mean squared displacement,
CRWs: correlated random walks,
LWs: Levy walks.

## Data sources

We provide position data from five separate experiments/movies on movements of Plasmodium-specific CD4 and CD8 T cells in murine brains that we generated from imaging data provided by the Bill Heath (lead author of the Ghazanfari *et al*. [27] paper). The data are provided as supplement to this paper (xlsx spreadsheet) and in independent repository https://github.com/DhruvPatel5701/TcellsInBrain. The data on movement of Toxoplasma-specific CD8 T cells were provided by Dr. Chris Hunter in 2013 and are available from Dr. Hunter upon request via.

## Codes

We provide several python codes with the publication. First, we provide the code that cleans cell trajectories; that is trajectories that have missing time frames are split to generate a new track. Second, we provide the code the calculate MSD and step/movement length distribution, and perform regression analysis on MSD data and tail analysis of movement length distribution. Third and finally, we provide a code to simulate movement of agents according to Brownian walk, Levy walk, or generalized Levy walk. The codes are available here: https://github.com/DhruvPatel5701/TcellsInBrain.

## Author’s contributions

VVG had the original idea of the study. VVG obtained position data from Harris et al. Nature 2013 paper. VVG also obtained imaging data from Ghazanfari et al. JI 2021 paper. RL segmented the imaging data using Imaris and generated coordinates for Plasmodium-specific CD4 and CD8 T cells in murine brains. Quality of segmentation was checked by VVG. BM performed initial analyses of the data. DP developed python codes and analyzed the data from both studies. DP also developed simulations for agents and analyzed the simulated data. BM and VVG supervised the work. DP and RL wrote the first draft of the paper, and VVG formulated the final draft. All authors contributed to the final version of the paper.

## Acknowledgements

This work was supported by the NIH grants (primarily by R01GM118553 and in part by R01AI158963) to VVG.

## Supplemental Information

### Additional details of tracing T cells using Imaris (ver 9.8.2)

The algorithm used for tracing the relevant movies was carefully designed, in terms of parameters, to maximize data retention and accuracy. The option of “Segment only a Region of Interest” was deselected as the desired region of interest was the entire image and not a segment. The option of “Process entire Image finally” was selected as the desired region of interest was the entire image. The option of “Different Spot Sizes (Region Growing)” was deselected as the region of interest indicated no signs of T-cell growth or development. The option of “Track Spots (over time)” was selected as this allowed each T-cell traced to have a unique set of output measurements. The option of “Classify Spots” was deselected as information on this feature was not provided by the manual and therefore left out of the algorithm. The option of “Object-Object Statistics” was deselected as the information provided by this option such as “Shortest Distance To Spots” and “Shortest Distance To Surfaces” was not needed nor did it affect the final tracing results. By deselecting this feature, this sped up the tracing process.

The “Source Channel” was changed to the channel containing the desired T-cells. The option of “Estimated XY Diameter” was set to 9.94 *µ*m. We found that a value of 9.94 was a safe value as Imaris detected nearly all of the desired T-cells. The option of “Model PSF-elongation along the Z-axis” was deselected as a safe value for the Z diameter was not determined in order to prevent the loss of T-cells. The option of “Background Subtraction” was selected as this feature smooths the final image. The filter type was set to “Quality” as this allowed for manual control of the threshold value for which Imaris detected a T-cell. The threshold value was set to a low value (less than 100) as this ensured that nearly all of the T-cells were detected. The next set of options under the “Create”, “Settings”, and “Color” tab are left alone as the editing of T-cells and T-cell tracks was done after the completion of the algorithm. Features under “Settings” and “Color” are personal preference and do not affect the final tracing results.

Afterwards, the “Tracking Algorithm” was set to “Autoregressive Motion” as this tracking algorithm best suited the data as the movement of T-cells were a continuous motion and not random. The next set of options were “Max Distance” and “Max Gap Size” which can be found under “Parameters”. These values were not tampered with as the values were automatically generated by Imaris and may vary from data set to data set. The option of “Fill gaps with all detected objects” was deselected to prevent the possibility of Imaris merging two independent T-cells into one T-cell resulting in error. The option of “Filter Type” was set to “Track Duration” as this allowed for manual control and the filtering of certain tracks with an “x” duration. The “Track Duration” value was set to a value less than 100 in order to prevent the loss of potential T-cell tracks. The option “Classification” and “Event Setup” were not added to the algorithm as information on this feature was not explained in the manual and therefore left out of the algorithm.

**Supplemental Movie S1:**
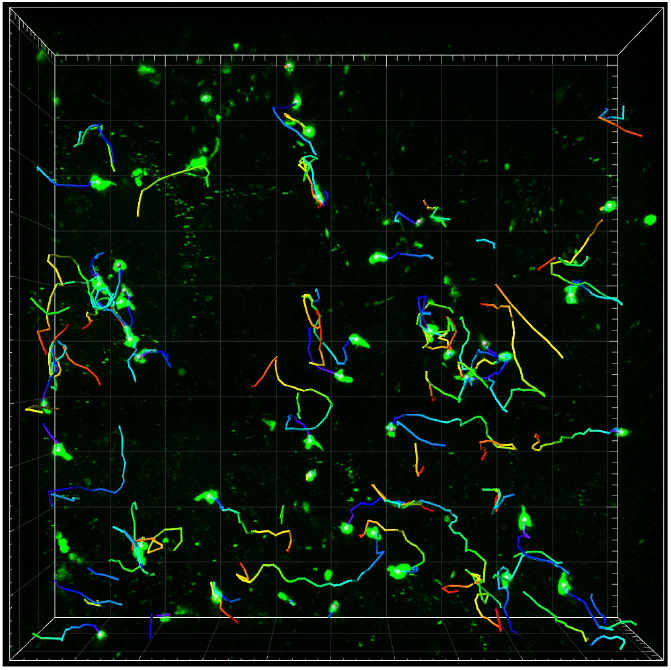
Example of movement of brain-localized Plasmoidum-specific CD4 and CD8 T cells at different times after *Plasmodium berghei ANKA* (PbA) infection. Experiments were performed as described in **Figure 1** in 5 individual mice. In the movie red are CD4 T cells and green are CD8 T cells. Bar scale is 50 *µ*m. The volume of 512 × 512 × 44 *µ*m was scanned approximately every 30 sec (exact imaging frequency varied by the movie, see Materials and methods for more detail).

**Supplemental Figure S1:**
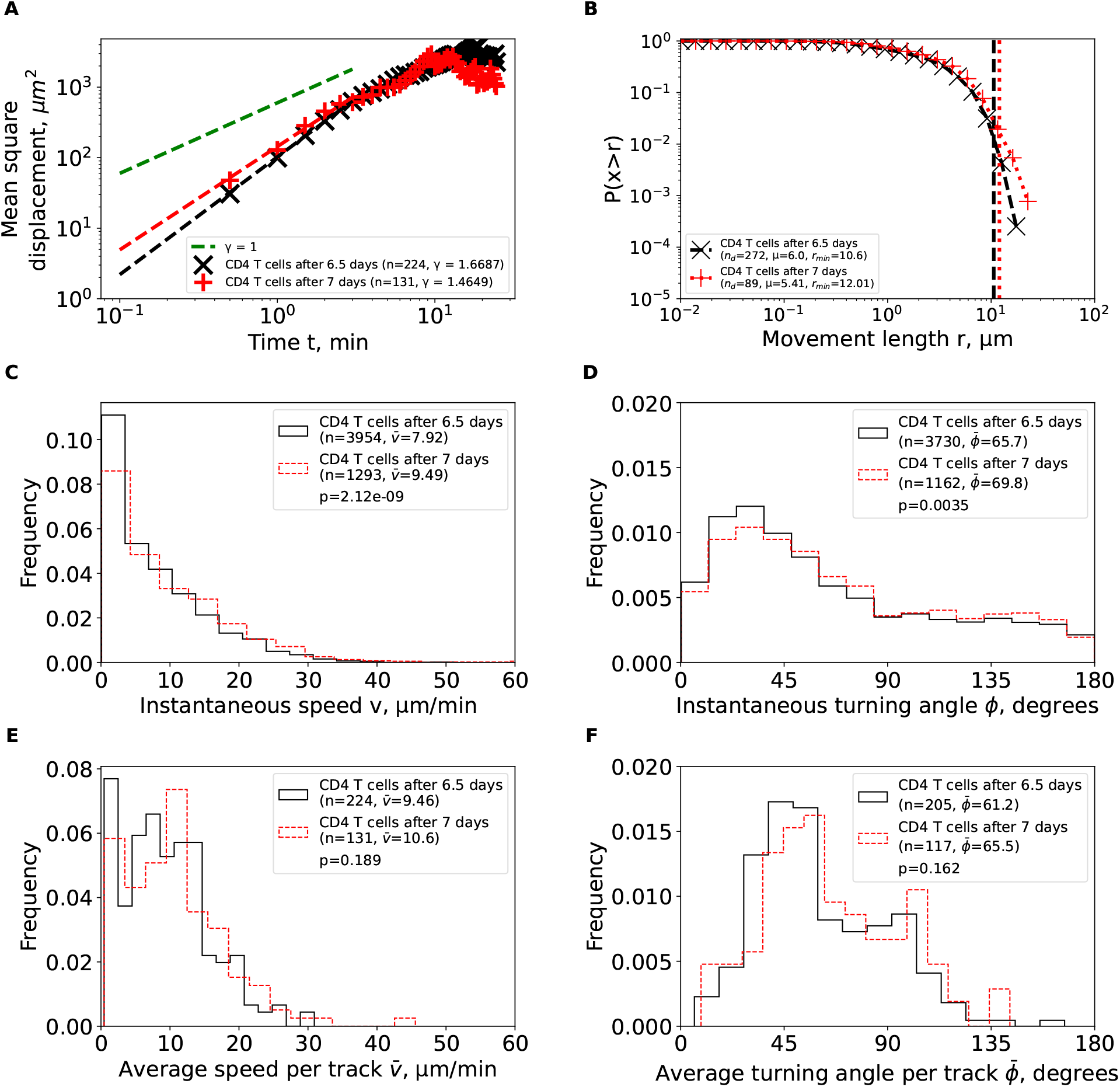
Brain-localized Plasmodium-specific CD4 T cells have higher instantaneous speed and higher turning angle at later times of PbA infection. The data from **Figure 3** we analyzed trajectories of CD4 T cells separately for 6.5 or 7 days since PbA infection. Other notations are the same as in **Figure 3**.

**Supplemental Figure S2:**
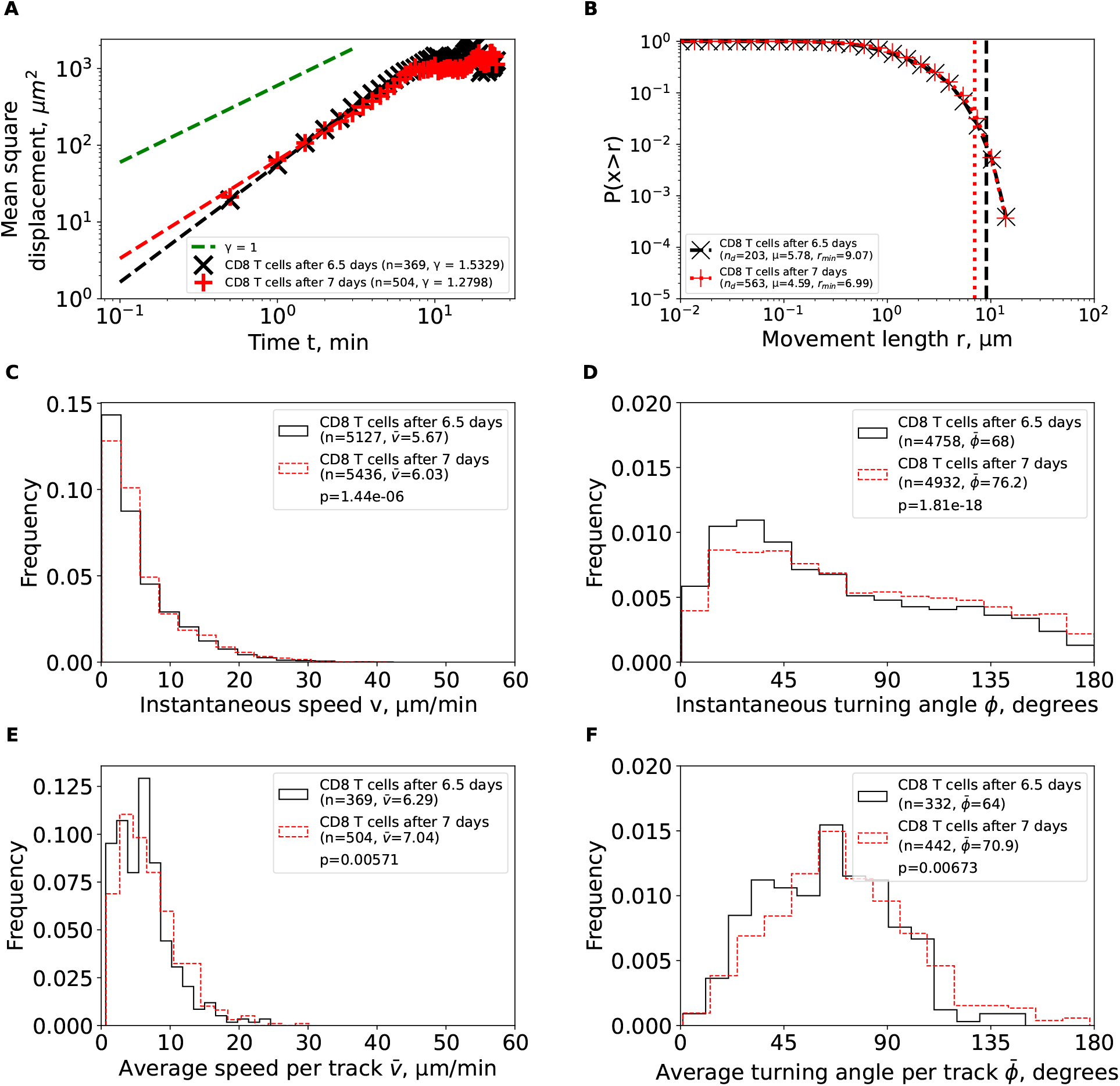
Brain-localized Plasmodium-specific CD8 T cells have higher speeds and higher turning angle at later times of PbA infection. The data from **Figure 3** we analyzed trajectories of CD8 T cells separately for 6.5 or 7 days since PbA infection. Other notations are the same as in **Figure 3**.

**Supplemental Figure S3:**
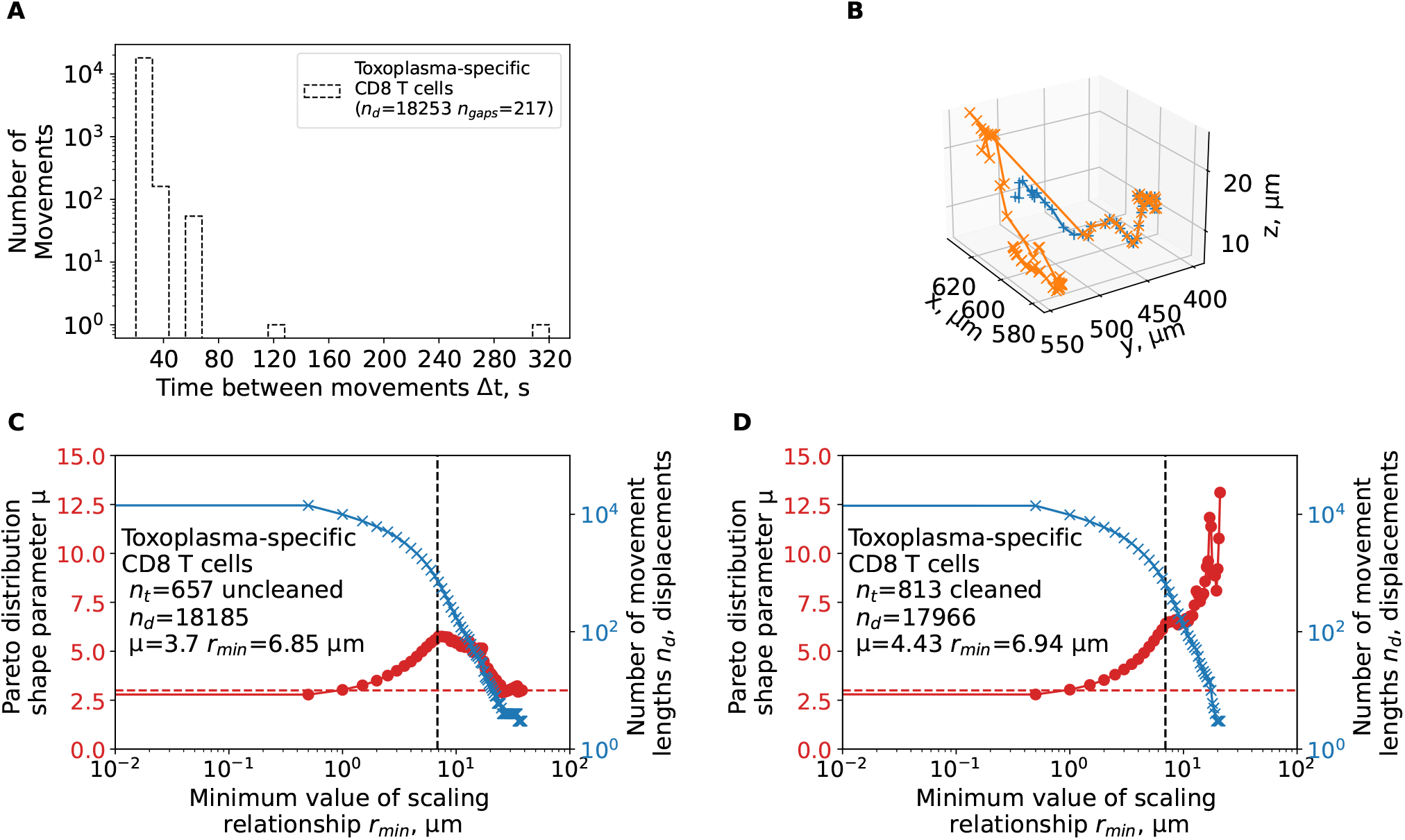
Artifacts in the track data have likely generated an appearance of Levy walks for brain-localized *Toxoplasma gondii* -specific CD8 T cells. We have reanalyzed track data from a previous publication [2] and calculated the shape parameter of the Pareto distribution using tail analysis of the movement length displacement data. **A**: The time between sequential movements in the data. **B**: a single large displacement for a trajectory yep.3829 observed in the data. **C-D**: We estimated the shape parameter *µ* of the Pareto distribution (eqn. (3), see Materials and Methods for details) by tail analysis of the movement length distributions for uncleaned (original) data (**C**) or when cell tracks were cleaned/split to allow for sequential timeframes for all trajectories (**D**). We also performed the analysis when we adjusted the tail cutoff value *r*_min_ (minimum value of the scale parameter) to different values denoted on the x-axis.

**Supplemental Figure S4:**
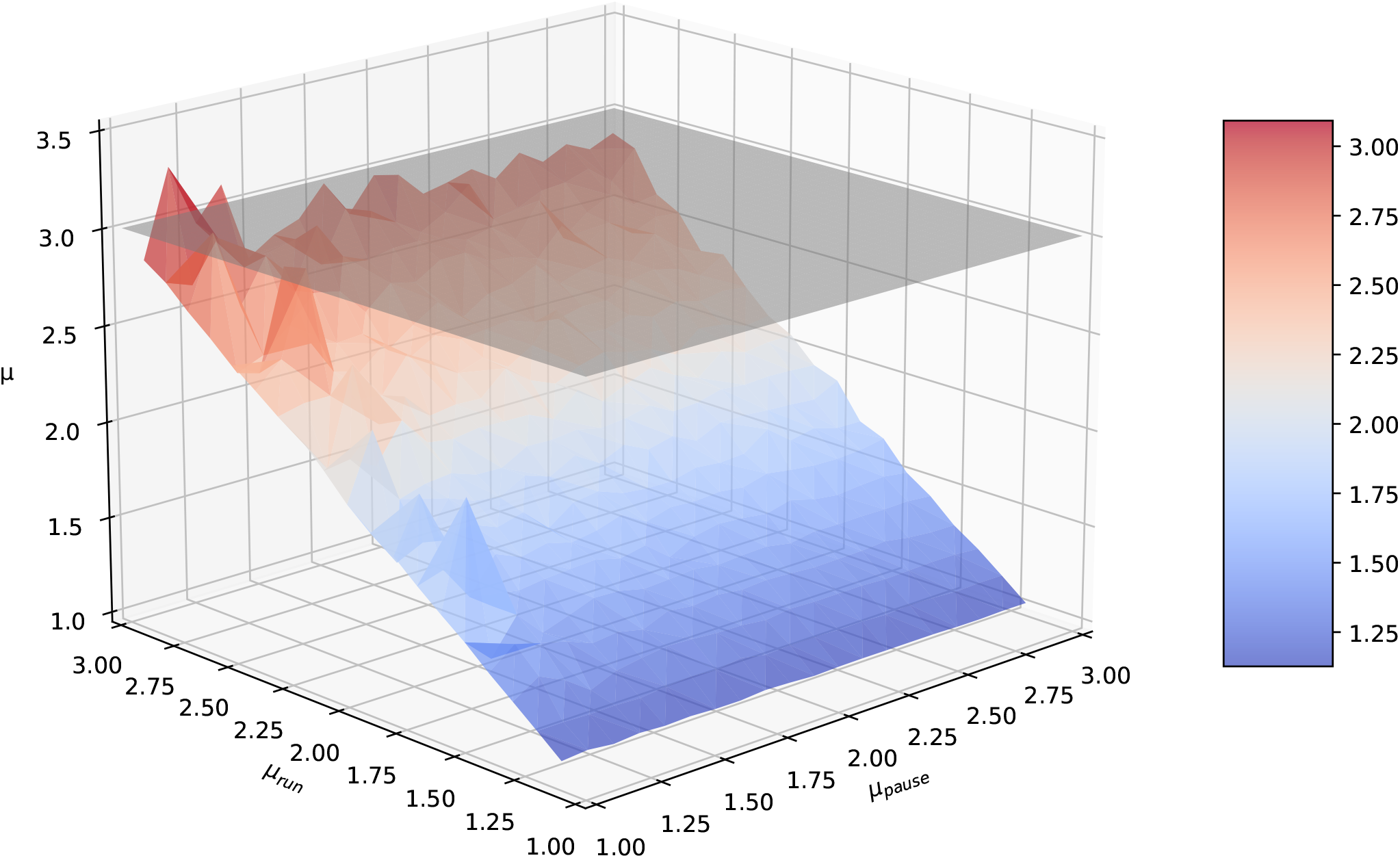
Tail analysis of movement lengths with python package powerlaw is a robust method to detect Levy walks. Generalized Levy walkers were generated for 1 < *µ*_*run*_ ≤ 3 and 1 ≤ *µ*_*pause*_ ≤ 3, in 0.1 increments per parameter with *r*_*min*_ = 1 for all Pareto distributions generated (see Materials and Methods for more detail). For each *µ*_*run*_ and *µ*_*pause*_ combination, 100 generalized Levy walkers were simulated for 100 steps. Then, the shape parameter *µ* of the tail of the movement length distribution was estimated using powerlaw package in python.

**Supplemental Table S1:** Estimated parameters of the generalized Pareto distribution fitted to the movement length data. We fitted the generalized Pareto distribution (eqn. (2)) using likelihood method (eqn. (4)) to different subsets of data including data from simulations. Using estimated parameters we also calculated the mean and variance of the distribution (see eqn. (2)). For every dataset we list the number of displacements used for fitting the GP distribution to the data, and “nan” stands for not-available. We typically simulated movements of 100 cells for 100 movements (with 10 sec for each movement).

| Cell population | # of displ | Shape $k$ | Location $\theta$ | Scale $\sigma$ | Mean | Variance |
| --- | --- | --- | --- | --- | --- | --- |
| All Plasmodium-specific CD4 T cells | 5257 | -0.08 | 0.01 | 4.47 | 4.15 | 14.84 |
| All Plasmodium-specific CD8 T cells | 10563 | -0.06 | 0.03 | 3.07 | 2.93 | 7.5 |
| Plasmodium-specific CD4 T cells at 6.5 days | 3954 | -0.09 | 0.04 | 4.28 | 3.97 | 13.03 |
| Plasmodium-specific CD4 T cells at 7 days | 1293 | -0.08 | 0.01 | 5.11 | 4.74 | 19.26 |
| Plasmodium-specific CD8 T cells at 6.5 days | 5127 | -0.04 | 0.03 | 2.93 | 2.84 | 7.24 |
| Plasmodium-specific CD8 T cells at 7 days | 5436 | -0.07 | 0.05 | 3.17 | 3.02 | 7.71 |
| All Toxoplasma-specific CD8 T cells | 18036 | 0.09 | 0.02 | 1.79 | 1.97 | 4.6 |
| *Simulated Brownian walkers* | 10000 | 0.15 | 1 | 0.27 | 1.32 | 0.14 |
| *Simulated CRW walkers* | 10000 | 0.15 | 1 | 0.27 | 1.32 | 0.14 |
| *Simulated Levy walkers* | 10000 | 0.72 | 1 | 0.65 | 3.32 | nan |
| *Simulated Generalized Levy walkers* | 1881 | 0.65 | 1 | 0.67 | 2.9 | nan |
| *Simulated Bullet motion walkers* | 10000 | 2.01 | 1 | 1.97 | ∞ | nan |

